# Predator-prey scaling laws support a suspension-feeding lifestyle in Cambrian luolishaniid lobopodians

**DOI:** 10.1101/2025.11.05.686786

**Authors:** Jared C. Richards, Javier Ortega-Hernández

**Affiliations:** Museum of Comparative Zoology and Department of Organismic and Evolutionary Biology, Harvard University, 26 Oxford Street, Cambridge, MA 02138, USA

**Keywords:** Luolishaniidae, Radiodonta, plankton, statistics, macroecology, niche ecology, functional morphology

## Abstract

The early Paleozoic saw a dramatic diversification of shelly epibenthic metazoans adapted to suspension and filter feeding, but the extent to which these radiations affected the evolution of non-biomineralized suspension-feeding taxa is uncertain because these organisms are not typically well represented in the fossil record. Luolishaniids are a highly derived and disparate clade of (typically) armoured lobopodians, widely interpreted as suspension feeders based on the presence of five or six anterior pairs of setulose appendages. Luolishaniids are globally widespread and represent the only Cambrian non-biomineralized free-living epibenthic bilaterians suggested to have a suspension-feeding mode of life, but their proposed ecology relies solely on a qualitative interpretation of their functional morphology. Here we test the hypothesis that the setulose appendages of luolishaniids were adapted for a suspension-feeding function. Quantitative morphological comparisons reveal a positive and statistically significant relationship between body length and the mesh spacing of the setulose anterior limbs of luolishaniids. Standardized comparisons indicate that the body size disparity between luolishaniids (predators) and Cambrian mesoplankton (prey) is consistent with patterns observed in modern suspension-feeding organisms. We provide quantitative evidence for suspension feeding in luolishaniids, which represents the first statistically supported example of modern-like predator-prey scaling patterns observed in Cambrian soft-bodied metazoans. Despite the uncanny appearance of luolishaniids, and Cambrian organisms more broadly, our results suggest their adaptations and mode of life feature ecological attributes shared with modern marine invertebrates.

## INTRODUCTION

Suspension feeding is a widespread life strategy in which animals capture food particles such as phytoplankton, zooplankton, and organic detritus from the water column in aquatic environments [1–3]. Suspension feeding is found throughout diverse animal clades, and may be further subcategorized based on precise habitat, mode of life, and locomotory capabilities of the organisms [1]. Given that the feeding strategy of these organisms fundamentally involves extracting organic matter from large volumes of water, they play a key ecological role as primary consumers in aquatic food webs where they help cycle nutrients, transport biomass from small particles into higher trophic levels in the water column, increase light penetration, and alter the distribution of planktonic lifeforms in space and time [1,4–7].

Suspension feeding among the earliest animal-dominated communities is well represented during the early Paleozoic among biomineralizing organisms with high preservation potential such as sponges, brachiopods, echinoderms, and bivalved mollusks [8–13]. By contrast, our understanding of the evolution of non-biomineralizing suspension-feeding organisms is more precarious due to their comparatively low preservation potential in the conventional fossil record [14–18] (Fig. 1). Radiodonts, typically large-bodied nektonic stem-group euarthropods typified by the presence of arthropodized frontal appendages, represent the most geographically widespread suspension-feeding soft-bodied metazoans found in sites with exceptional preservation around the world during the early Paleozoic [19,20]. Suspension-feeding radiodonts include the hurdiids, which become increasingly diverse after the middle Cambrian and extended into the Devonian [21–25,25,26] and the rarer tamisiocaridids (Fig. 1C), which are restricted to the early Cambrian [27–29]. Other Cambrian soft-bodied organisms with a likely suspension-feeding mode of life based on comparisons with modern analogues include cnidarians, hemichordates, tunicates, vertebrates, ctenophores, and bivalved euarthropods [30–35] (Fig. 1D, 1E). Suspension feeding has also been proposed for luolishaniids, a diverse group of lobopodians found in several major Cambrian sites with exceptional preservation around the world [36–43] (Fig. 1A, B). The general body plan of luolishaniids consists of a tubular trunk with five or six pairs of differentiated setulose anterior appendages, followed by a variable number of clawed walking legs on the trunk. Other aspects of their morphology vary substantially, such as extent of spinose dorsal amour ranging from absent (*e.g.*, *Facivermis, Ovatiovermis*) to extravagant (*e.g.*, *Collinsium, Collinsovermis, Acinocricus*), as well as their body size (roughly 10 to 100 mm in length) and lifestyle (*e.g.*, free living or tube dwelling) [39,42]. Luolishaniids differ from other suspension-feeding organisms because most of them had a vagile epibenthic mode of life and lack obvious extant analogues, with phylogenetic analyses suggesting affinities as either early relatives of velvet worms (Onychophora), water bears (Tardigrada), or even panarthropods more broadly [39,42,44,45]. However, the view of luolishaniids as suspension feeders is exclusively based on the qualitative interpretation of the functional morphology of their anterior setulose appendages, and thus this ecological hypothesis remains to be critically tested, especially considering the peculiar morphology and variable lifestyle of these organisms. Other possible functional interpretations for the prominent setulose anterior appendages of luolishaniids include grasping macroscopic prey through raptorial-like feeding, or possibly to sieve the sediment in search for food items as proposed for some radiodonts [46,47].

**Figure 1.**
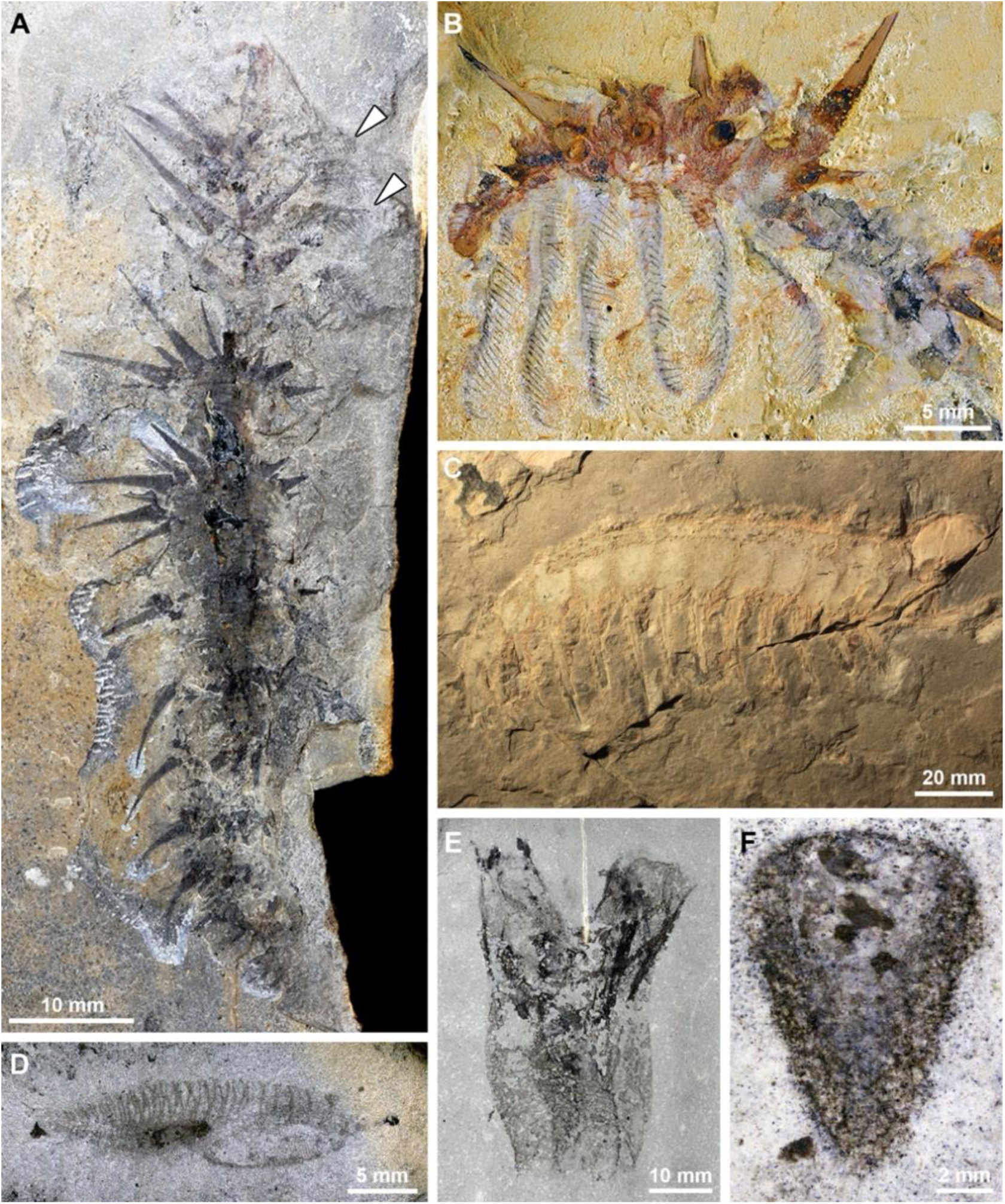
Diversity of exceptionally preserved suspension feeding organisms from Cambrian deposits. **A)** The luolishaniid lobopodian *Acinocricus stichus* [41] from the middle Cambrian (Wuluian) Spence Shale, Utah (Kansas University Museum Invertebrate Paleontology collection; KUMIP 204353); arrowheads indicate position of anterior setulose appendages. **B**) The luolishaniid lobopodian *Collinsium ciliosum* [39] from the early Cambrian (Stage 3) Xiasohiba biota, South China (Yunnan Key Laboratory for Paleobiology; YKLP 12127), showing well preserved six pairs of anterior setulose appendages. **C)** Frontal appendage of the tamisiocaridid radiodont *Echidnacaris briggsi* [29] from the early Cambrian (Stage 4) Emu Bay shale, South Australia (South Australian Museum, Adelaide Paleontological collection; SAMA P40180). **D)** The vertebrate *Nuucicthys rhynchocephalus* [34] from the mid-Cambrian (Drumian) Marjum Formation, Utah (Utah Museum of Natural History; UMNH.IP.6084). **E)** The tunicate *Megasiphon thylakos* [31] from the mid-Cambrian (Drumian) Marjum Formation, Utah (UMNH.IP.6079). **F.** The hexactinellid-like sponge *Polygoniella turrelli* [13] from the mid-Cambrian (Drumian) Marjum Formation, Utah (Museum of Comparative Zoology Invertebrate Paleontology collection, Harvard University; MCZ.IP.199049).

In this study we quantitatively test the hypothesis that the functional morphology of Cambrian luolishaniids is congruent with a suspension-feeding mode of life by examining the relationship between body size and sieving potential of the setulose anterior appendages. Based on comparative statistical methods, we provide the first quantitative evidence for suspension feeding in luolishaniids and demonstrate that these lobopodians follow the same suspension feeder to prey size scaling laws observed in modern ecosystems.

## MATERIAL AND METHODS

### Data collection

High-resolution digital images from all previously described luolishaniid taxa were loaded into Adobe Illustrator 28.2 (2024) (see Table 1). We measured the spacing between the setae (mesh spacing) on the anterior limbs for each taxon, as this is generally considered a proxy for determining the size of seston collected by suspension feeders [27,48,49]. Given the paucity of complete specimens and the widespread likelihood of morphological distortion via diagenesis, we used the maximum recorded trunk length and width in the original description (Table 1). In cases where the specimen with the best preservation of the setulose structures is also complete, we would collect the body size measurement from the same specimens.

**Table 1.** Autecological metrics for all Cambrian luolishaniid lobopodians. Note that trunk length in the Emu Bay Shale luolishaniid (asterisk) is estimated from the statistically significant scaling pattern established amongst other Cambrian luolishaniids.

| Taxa | Number of Mesh Spacings (N) | Mean Mesh Spacing ($\mu\text{m}$) | Minimum Estimated Prey Diameter ($\mu\text{m}$) | Maximum Estimated Prey Diameter ($\mu\text{m}$) | Trunk Length (mm) | Trunk Width (mm) |
| --- | --- | --- | --- | --- | --- | --- |
| *Acinocricus stichus* [41] | 21 | 319.3 | 457.1 | 4248.5 | 96.0 | 13.2 |
| *Collinsium ciliosum* [39] | 52 | 250.6 | 358.0 | 3421.5 | 85.0 | 4.2 |
| *Collinsovermis monstrosus* [41] | 25 | 206.2 | 294.1 | 2875.1 | 35.0 | 7.6 |
| *Enthothyreos synnaustrus* [43] | 14 | 290.4 | 415.4 | 3902.8 | 38.12 | 5.2 |
| *Facivermis yunnanicus* [42] | 8 | 202.7 | 289.1 | 2831.3 | 90.0 | 2 |
| *Luolishania longicruris* [37] | 8 | 108.4 | 153.8 | 1619.4 | 14.30 | 1 |
| *Ovatiovermis cribratus* [40] | 26 | 161.1 | 229.3 | 2305.8 | 18.0 | 3.3 |
| Collins' monster, Emu Bay [38] | 12 | 272.0 | 389.0 | 3681.6 | 87.5* | - |

**Table 2.** Predator: Prey ESD Ratios by Clade. Luolishaniid body sizes calculated by modelling taxa as cyliders (*V* = π*r*^2^*h*) to obtain their volume and converting to ESD for standardization.

| Clade | N | Mean Ratio | Median Ratio | $\log_{10}(\text{Mean Ratio})$ | $\log_{10}(\text{Median Ratio})$ |
| --- | --- | --- | --- | --- | --- |
| Chaetognatha | 6 | 4.9 | 5.0 | 0.7 | 0.7 |
| Cladocera | 5 | 45.4 | 47.2 | 1.7 | 1.7 |
| Copepoda | 9 | 20.5 | 17.5 | 1.3 | 1.2 |
| Luolishaniidae | 7 | 36.1 | 29.0 | 1.5 | 1.5 |
| Rotifera | 4 | 17.3 | 17.3 | 1.2 | 1.2 |
| Trochozoa | 7 | 39.9 | 24.4 | 1.4 | 1.4 |

### Data Analysis

All analysis was completed in RStudio environment using the packages dplyr, plyr, magrittr, stats, lm4, lmerTest, performance, MuMIn, lmtest, ggrepel, and ggplot2 [50–60].

Given the smaller sample size of the available data (*e.g.,* multiple mesh spacing values imposed onto only one trunk length measurement per species), we used a variety of analytical tools to extract as much information as possible from our limited dataset while being methodologically conservative. We employed an Ordinary Least-Squares (OLS) linear mixed-effect model with a gaussian response distribution and an identity link function with Satterthwaite’s approximation to explore the relationship between genus-level body length and unaggregated mesh spacing. We treat genus-level body length (mm) as the fixed effect and unaggregated mesh spacing as the dependent variable (*µ*m). To account for the nested structure of the data, we included genus as a random effect in the model. This model was used to determine whether there was a statistically significant relationship between luolishaniid genus body length and corresponding mesh size while also controlling for non-independent observations [48,49,54,61]. Next, we performed an OLS linear regression examining the relationship between mean (aggregated) mesh size values and body size. This summarized view of the mesh spacing data would be used later for further examination of the suspension-feeding ecology of each luolishaniid species. We report the F-statistic, the coefficient of determination (R^2^), and the *p*-value recovered from the t-test of the regression’s slope coefficient (H_0_: regression slope = 0) for both OLS models. We also ran a Harvey-Collier test on the aggregated mesh spacing dataset to check the assumption of a linear relationship between body and mesh size on the logarithmic scale (H_0_: regression is correctly modeled as linear). Finally, using the relationship that exists between mesh spacing and estimated prey size for a variety of extant metazoans [27] and the mean mesh spacing values for each luolishaniid genus, we calculated the minimum and maximum estimated prey size (*µ*m) of all known luolishaniids to examine the lifestyles of this rare clade even further.

Neontological studies frequently use the equivalent spherical distance (ESD) to represent the size of suspensivores and their prey [49,62,63]. This measure allows the size of irregularly shaped objects, such as organisms, to be standardized into a 1-dimensional measure, in this case the diameter of a perfect sphere. We modelled our luolishaniids as cylinders (*V* = π*r*^2^*h*), where *h* and 2*r* represent the trunk length and width, respectively. This cylindrical volume was then converted into ESD using the formula ESD = (6*r*^2^*h*)^1/3^ [64]. The minimum estimated prey sizes previously calculated for luolishaniids were used as a proxy for their corresponding prey ESD, as this represents a more conservative value and has been shown to be more strongly correlated with mesh spacing in extant metazoans [27]. To determine if the predator-prey size relationships of luolishaniids were in line with those of known suspension-feeding clades, the modelled data for Cambrian luolishaniids was appended onto previously compiled data on predator and prey ESDs for modern suspension-feeding invertebrates collected from the literature [49,62,63]. Chaetognaths, a phylum of obligate ambush raptorial predators of similar size to luolishaniids that do not engage in suspension feeding [65,66] were included in previous studies of predator-prey relationships on the ESD scale in marine invertebrates. Given that they can be used as benchmark for the predator prey size ratios seen in active, marine invertebrate hunters, we included them in our study for comparison with luolishaniids and modern suspension feeders, but do not include them in the OLS analysis.

We performed an OLS linear regression on the logarithmic scale to determine if the relationships between predator and prey ESD between luolishaniids and modern suspension-feeding invertebrates was similar. We report the F-statistic, the coefficient of determination (R^2^), and the *p*-value recovered from the t-test of the regression’s slope coefficient (H_0_: regression slope = 0) of the OLS model. We also ran a Harvey-Collier test on the data to check that the linearity assumption was met.

## RESULTS

### Luolishaniid Allometry and Prey Size Estimate

We find a positive and statistically significant relationship between body size (mm) and mesh spacing (*µ*m) on the logarithmic scale for both unaggregated mesh spacing data (*F*(1, 4.82) = 8.52, *p*-value = 3.45×10^-2^, marginal R^2^ = 0.35, conditional R^2^ = 0.65) and aggregated mean mesh spacing data (*F*(1,5) = 7.72, *p*-value= 3.91×10^-2^, R^2^ = 0.61; S1_OLS; Fig. 2A). The corresponding sublinear power functions are Log_10_(Ŷ) = 0.36 × Log_10_(X) + 1.72 and Log_10_(Ŷ) = 0.37 × Log_10_(X) + 1.72. The linearity assumption was met in the OLS of aggregated mesh spacing data on the logarithmic scale (Harvey-Collier statistic = 0.36, degrees of freedom = 4, *p*-value = 0.74).

**Figure 2.**
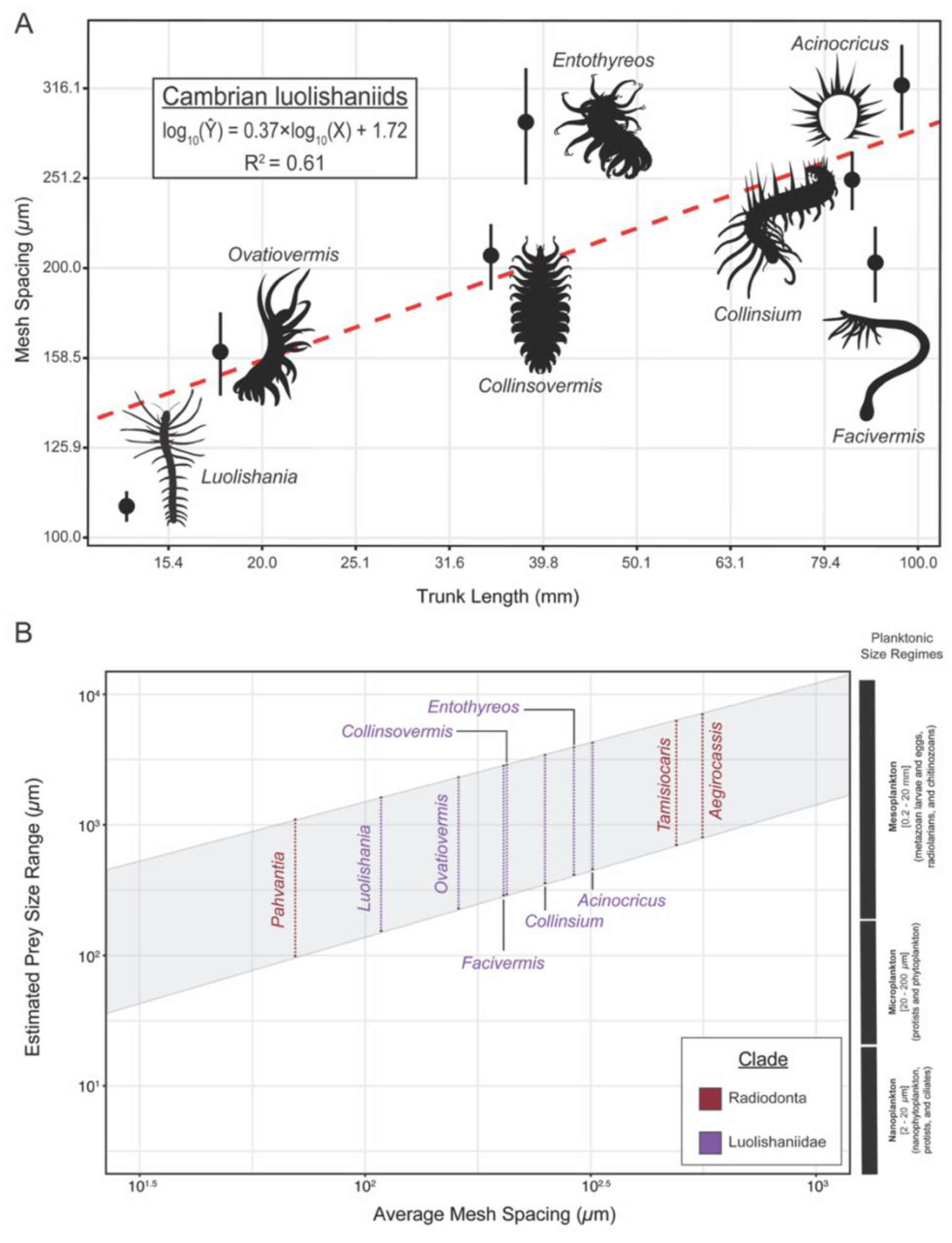
Quantitative comparisons between Cambrian suspension feeding soft-bodied metazoans. A) Allometry of mesh size of Cambrian luolishaniids. Relationship between body length (mm) and appendage mesh spacing (μm) among Cambrian luolishaniids, showing a significant positive trend (red dashed line; Log_10_(Ŷ) = 0.37 • Log_10_(X) + 1.72, R² = 0.61). Each point represents a species with a 95% CI around the mean mesh spacing. Silhouettes illustrate representative morphologies for each taxon; note that *Acinocricus* is represented by an isolated sclerite ring. B) Comparison between mesh spacing in luolishaniids (purple) and radiodonts (red) against estimated prey size. Radiodont data based on refs [23,27].

We find that the estimated minimum particle diameters would have enabled most luolishaniids to capture suspended prey and particles on the lower end of the mesoplanktonic scale (∼230 – 460 *µ*m) (Fig. 2B; Table 1). *Luolishania* is the only taxon estimated to have been able to collect suspended prey and particles that were on the microplanktonic spectrum (∼150 *µ*m) (Fig. 2B; Table 1). Given that the unnamed Emu Bay luolishaniid is based on a single incomplete specimen that obstructs our ability to measure it [38], we used the previously derived power laws to calculate its estimated maximum body size at approximately 87.5 mm (Table 1). However, due to the incompleteness of the only known specimen we refrain from including it in our main analyses to minimize potential sources of error.

### Luolishaniid Macroecology and Modern Suspension feeders

We find a positive and statistically significant relationship between predator and prey ESD on the logarithmic scale for analysis of luolishaniids and modern suspension-feeding invertebrate groups (*F*(1,30) = 198.1, *p*-value = 9.40×10^-15^, R^2^ = 0.87). The corresponding sublinear power function is Log_10_(Ŷ) = 0.90 × Log_10_(X) - 1.13. (Fig. 3A). The linearity assumption on the logarithmic scale was met in the OLS (Harvey-Collier statistic = 0.91, degrees of freedom = 29, *p*-value = 0.37), respectively.

**Figure 3.**
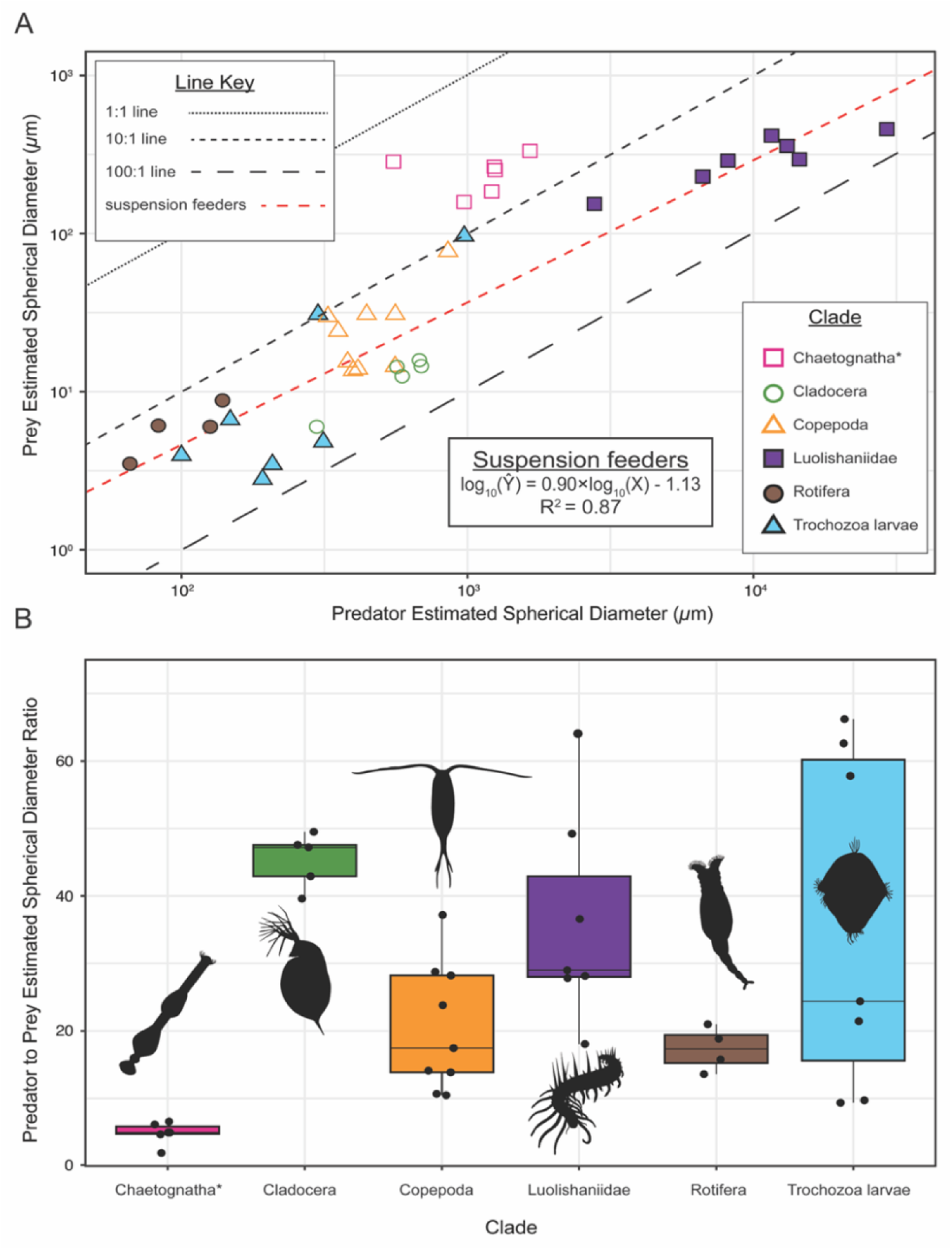
Quantitative comparisons between predator-prey body size scaling. A) Relationship between predator and prey size in extant and extinct marine invertebrates. Plot of estimated spherical distance (*μ*m) of marine invertebrates and their planktonic prey, showing a significant positive trend (red dashed line; Log_10_(Ŷ) = 0.90 • Log_10_(X) - 1.13, R² = 0.87). Note that chaetognaths represent obligate raptorial hunters, so they are shown for comparison with the remaining suspension-feeding invertebrate groups and were not included in the linear regression line. B) Predator-Prey ESD Ratios for multiple marine invertebrate clades.

There is a considerable range in the predator-prey ESD ratios between clades (Figure 3B). The group most similar in size to their prey are chaetognaths, which are on average five times larger. Copepods and rotifers are approximately 20 times larger than their prey. The ratio amongst luolishaniids, trochozoans, and cladocerans ranges approximately between 25–50, indicating that most suspension-feeding clades are between one to two orders of magnitude larger than their prey on the ESD scale. The differences in the predator-prey ESD ratios observed in confirmed suspension feeding extant organisms support this mode of life for Cambrian luolishaniids and allow us to reject the alternative hypothesis that these lobopodians raptorial feeders.

## DISCUSSION

### Luolishaniid Macroecological Scaling and Estimated Prey Size

The positive, statistically significant relationship between trunk length and mesh spacing in the setulose appendages of Cambrian luolishaniids (Figure 2A, B) on the logarithmic scale indicates that mesh spacing does not scale uniformly with body size. For a given unit of increase in body size, there is a proportionally smaller increase in mesh spacing. These observations recapitulate sublinear allometric laws of body size in extant animals [67], particularly among suspension-feeding aquatic invertebrates [48,68,69]. The strong (R^2^ = 0.87, *p*-value < 0.001) relationship between predator and prey ESD on the logarithmic scale (Figure 3A), provides quantitative support to the hypothesis that Cambrian luolishaniids were suspension feeders. The setulose appendages of luolishaniids would enable them to collect seston approximately 30–35 times smaller than their body size (when expressed in terms of ESD scaling), mirroring the predator-prey size relationship seen in predominantly filter-feeding groups such as larval mollusks and cladocerans [70–73] (Figure 3B). In contrast, obligate small invertebrate raptorial predators such as chaetognaths use their elongate anterior spines to actively ambush and seize relatively large prey when compared to their own body size [65,66].

In the context of luolishaniid feeding ecology, mesh size would have had direct implications for the planktonic prey targeted by the group. It is impossible to evaluate the precise composition of luolishaniid diets because all their fossils to date lack evidence of gut contents or other indications of food selectivity; thus, the mesh size provides the best available estimate of their prey items. We find that the smallest Cambrian luolishaniids had the finest mesh size, estimated at 108.4 *µ*m for *Luolishania* (body length = 14 mm) and 161.1 *µ*m for *Ovatiovermis* (body length = 18 mm) (Figure 2A, Table 1), which would have allowed them to capture larger microplankton (*e.g.*, phytoplankton), as well as the smaller cohorts of mesoplankton. All other larger-bodied luolishaniids, ranging from *Collinsovermis* (body length = 35 mm) to *Acinocricus* (body length = 96 mm), show a progressive increase in the mesh size on the setulose appendages, ranging from ca. 200 *µ*m to 450 *µ*m. This mesh size range would have allowed medium to large bodied luolishaniids to feed on all types of mesoplankton but would be less suitable for targeting microplankton as it would fall below the minimum capturable size (Figure 2B). More broadly, the range of estimated prey sizes of luolishaniids overlaps with that of suspension-feeding radiodonts such as *Aegirocassis benmoulai, Pahvantia hastata*, and *Tamisiocaris borealis* [20,23,27]. These comparisons further support the hypothesis of luolishaniids as suspension feeders, as both groups follow similar patterns of mesh spacing on their feeding appendages despite the substantial disparity in overall body size between the smaller luolishaniids (Table 1) and the typically much larger radiodonts (*A. benmoulai*, ca. 200 cm trunk length; *P. hastana,* ca. 24 cm estimated trunk length; *T. borealis,* ca. 34 cm estimated trunk length) [20,23,27]. These observations suggest that mesoplankton and larger microplankton were most likely the preferred food sources of multiple suspension-feeding clades during the Cambrian and the Early Ordovician. Critically, luolishaniids provide a valuable and previously unavailable insight into the relationships between body size and prey size (estimated by mesh size) of Cambrian non-biomineralized metazoans because of the availability of complete body fossils in several members of the same clade. By contrast, this information is absent from suspension-feeding radiodonts due to the paucity of complete body fossils in the Cambrian record.

The microfossil record of planktonic life during the early Paleozoic offers insights into the likely prey items that would have served as the primary food source of luolishaniids. Although the diversity and abundance of phytoplankton (*e.g.*, cyanobacteria, acritarchs) increased rapidly throughout the Cambrian, these microorganisms typically have body sizes below 100 *μ*m in diameter [74–78]. This range indicates that Cambrian phytoplankton would likely be too small for most luolishaniids based on their mesh spacing; only the smallest luolishaniids, namely *Luolishania* and *Ovatiovermis*, would have been able to feed off the largest phytoplankton (Figure 2B). Most luolishaniids would have been instead well suited for feeding on mesoplankton (ca. 0.2 to 2 mm diameter), which would include diverse organisms such as chitinozoans, radiolarians, and various types of metazoan larvae (i.e. zooplankton). Fossil evidence for chitinozoans dates to mid-Cambrian strata [79], although they are more common from the Ordovician to Devonian [80,81]. Chitinozoans likely extended into the early Cambrian, but it is possible that they were less abundant and thus did not produce a robust fossil record at the time. Thus, we posit that zooplankton likely represented the primary prey source for most Cambrian luolishaniids, which would have included the swimming larvae of cnidarians, arthropods, and spiralians such as brachiopods and mollusks [9,82–95].

### Comparisons to Modern Analogs

Luolishaniid lobopodians are unusual among early metazoans because they are largely soft-bodied (except for their sclerotized dorsal spines), suspension-feeding, epibenthic, vagile, and typically free living (with the possible exception of *Facivermis* [42]), and thus occupied a specialized ecological niche in Cambrian ecosystems. It seems plausible that other Cambrian metazoans also fit some of these attributes, particularly when considering the substantial diversity of filter-feeding apparatuses recovered from small carbonaceous fossils around the world [96–98]. For example, the early Cambrian annelids *Dannychaeta* [99] and *Iotuba* [100] are likely candidates of additional soft-bodied suspension-feeding metazoans based on comparisons with some living megaloniid and acrociriid polychaetes. However, further discoveries of complete organisms with well-preserved feeding structures are needed to confirm this possibility. Luolishaniids have been directly compared with some extant suspension-feeding invertebrates, particularly caprellid amphipod crustaceans, based on suggested similarities in their morphology including an elongate tubular body, anterior setulose appendages with a sieving function, and clawed walking legs on body trunk or posterior for grasping onto the substrate [40,41,101–103]. However, these taxa differ in important aspects of their functional morphology with direct implications for their precise mode of life and feeding strategies. Caprellid amphipods employ their cilia-laden, mucus-covered antennae to initiate suspension feeding [40,101]. After detecting a substantial accumulation of seston on the antennae, caprellids use their raptorial gnathopods to cleanse these appendages from food particles, form a ball of mucus and seston, and pass it to their mouth for consumption [101].

Although luolishaniids and caprellid amphipods employ setulose anterior appendages to capture plankton, the former differs in the absence of specialized cleaning limbs or complex mouthparts employed in suspension feeding [39,40,42,101,102,104]. These differences suggest that luolishaniids most likely employed a different strategy to consume seston, as there is no evidence whatsoever that luolishaniids could secrete or employ mucus for feeding. Instead, suction feeding would likely be a suitable option as it is widely employed by aquatic invertebrates where a negative pressure vacuum is produced to draw nearby water, and therefore any suspended materials, into the buccal cavity [105,106]. This interpretation is in line with the recent hypothesis that armored lobopodians were adapted for suction feeding based on the presence of a muscular pharynx (e.g. *Onychodictyon ferox*) and aciculate sclerotized teeth in the foregut with a proposed filtering function in the hallucigeniid *Hallucgenia sparsa* and the luolishaniid *Ovatiovermis cribatus* [107]. Experimental work on the fluid dynamics of extant suction feeders has also shown that no modern invertebrates with hydrostatically supported bodies can create the same suction forces seen in aquatic vertebrates such as fish [108]. Therefore, we hypothesize that the suction feeding capacity of luolishaniids would have been comparatively low, just enough so that a low-pressure difference between the buccal cavity and the environment would suffice to consume the prey captured by the sieving apparatus on their anterior setulose appendages.

### Broader Paleoecological Implications

Luolishaniids differ from other Cambrian suspension-feeding soft-bodied animals, particularly radiodonts such as *Tamisiocaris*, *Pahvantia,* or *Hurdia*, in that the former lobopodians most likely had a slow-moving epibenthic habitus, which means they would not be able to force water over their sieving apparatus by continuously moving against the water column [20,27]. Instead, luolishaniids most likely sought after microhabitats with low to intermediate water flow rates facing their anterior setulose limbs perpendicular to the oncoming water current to collect suspended food items [1,109–111]. The water flow speeds necessary for stationary suspension feeding of mesoplankton in modern ecosystems range from 5 to 10 cm/s based on experiments involving modern suspension feeders feeding on mesoplankton [110,112]. Assuming that luolishaniids would require a similar flow speed necessary for effective suspension feeding, the role of the robust sclerotized terminal claws on the posterior trunk appendages becomes critical, as it would allow them to maintain a secure attachment to hard substrates during food capture without becoming dislodged [1,39,110,112,113].

The feeding ecology of luolishaniids might also have implications for better understanding their mode of life considering their peculiar morphology, particularly the presence of extensive dorsal spinose defensive armature in most species [39,41,43]. Neontological behavioural work has shown that free living animals that bear some type of external body armor (*e.g.*, epidermal spines, plates or scales) display a higher threat tolerance or threshold before they flee or take refuge from potential predators when compared to unarmored forms, both in aquatic and terrestrial environments [114,115]. Applied in the context of luolishaniid paleobiology, this generalized behavior would suggest that smaller-bodied species and/or those in which dorsal armor is relatively light (*Luolishania*) or non-existent (*Ovatiovermis*) would be more hasty to seek refuge compared to larger bodied and more heavily armored forms, such as *Acinocricus, Collinsium, Collinsovermis,* and *Entothyreos* [38,39,43]. *Facivermis* is a notable exception in this regard because it is large-bodied (ca. 90 mm long) but completely lacks dorsal armor [42]. In this case, the unique tube-dwelling mode of life of *Facivermis* most likely represents a tradeoff that allowed it to attain a large body size despite the absence of dorsal armor by shifting to a different defensive strategy that instead relies on the protection of the secreted tube, facilitating a readily accessible refuge if disturbed. It is notable that the extent of dorsal armature in luolishaniids closely follows their overall body size, with *Acinocricus, Collinsium, Collinsovermis, Entothyreos* featuring the heaviest coverage within the clade based on the number of dorsal spines per appendage pair. The heavy armature of these forms would have likely made their movements cumbersome due to excessive resistance against the water column and thus caused a tradeoff against their ability to swiftly escape from potential predators, as also observed in modern species. Heavily armored luolishaniids would have primarily relied on their body size and impressive arrays of dorsal spines as primary deterrents, particularly when actively engaging in suspension feeding against the water current [114–117].

In conclusion, our results provide novel quantitative support to the longstanding interpretation of luolishaniids as epibenthic suspension feeders and suggest that they follow comparable macroecologial patterns observed in modern invertebrate clades with a suspension feeding autecology. The findings indicate that the fundamental requirements for a suspension-feeding lifestyle among non-biomineralized metazoans were already in place by the early and middle Cambrian based on a comparative quantification of luolishaniid morphology. This study also provides the first estimates of planktonic prey sizes for luolishaniids and provides further novel insights into their likely behavior based on modern analogues. Luolishaniids are among the most otherworldly animals that evolved during the Cambrian Explosion, hence their initial moniker as “Collin’s monsters”, honoring Desmond Collins who illustrated the first example of these bizarre organisms in 1986 from a fossil collection produced by the Royal Ontario Museum field party in 1983 [38,41]. Instead, we posit that main adaptations and mode of life of luolishaniids can be readily explained through the lens of a modern marine suspension-feeding ecology.

## ACKNOWLEDGEMENTS

We thank Rudy Lerosey-Aubril (Harvard University) for sharing digital images of Marjum fossils, and John Paterson (University of New England) for sharing the digital image of *Echidnacaris briggsi*. Thanks to Yang Jie and Xi-guang Zhang (Yunnan University) for sharing the digital image of *Colllinsium.* Thanks to Bruce Lieberman (The University of Kansas) for facilitating access to the holotype of *Acinocricus stichus*. This work is funded by NSF CAREER award no. 2047192 “Ecological Turnover at the Dawn of the Great Ordovician Biodiversification Event—Quantifying the Cambro-Ordovician Transition through the Lens of Exceptional Preservation,” and NSF GRFP award no. 2140743.

